# Reproductive workers show queen-like gene expression in an intermediately eusocial insect, the buff-tailed bumble bee *Bombus terrestris*

**DOI:** 10.1101/012500

**Authors:** MC Harrison, RL Hammond, EB Mallon

## Abstract

Bumble bees represent a taxon with an intermediate level of eusociality within Hymenoptera. The clear division of reproduction between a single founding queen and the largely sterile workers is characteristic for highly eusocial species, whereas the morphological similarity between the bumble bee queen and the workers is typical for more primitively eusocial hymenopterans. Also, unlike other highly eusocial hymenopterans, division of labour among worker sub-castes is plastic and not predetermined by morphology or age. We conducted a differential expression analysis based on RNA-seq data from 11 combinations of developmental stage and caste to investigate how a single genome can produce the distinct castes of queens, workers and males in the buff-tailed bumble bee *Bombus terrestris*. Based on expression patterns, we found males to be the most distinct of all adult castes (2,411 transcripts differentially expressed compared to non-reproductive workers). However, only relatively few transcripts were differentially expressed between males and workers during development (larvae: 71, pupae: 162). This indicates the need for more distinct expression patterns to control behaviour and physiology in adults compared to those required to create different morphologies. Among female castes, reproductive workers and their non-reproductive sisters displayed differential expression in over ten times more transcripts compared to the differential expression found between reproductive workers and their mother queen. This suggests a strong shift towards a more queen-like behaviour and physiology when a worker becomes fertile. This contrasts with eusocial species where reproductive workers are more similar to non-reproductive workers than the queen.

## Introduction

Eusociality, the division of adult females in reproductive queens and mainly sterile workers that care for the brood, has evolved multiple times independently within the Hymenoptera (bees, ants and wasps; Andersson 1984). The level of sociality varies within the Hymenoptera, ranging from non-social solitary species through primitively eusocial to highly eusocial taxa. Among highly eusocial hymenopterans, beside the clear division of reproduction between morphologically distinct workers and queens, further worker sub-castes exist. These worker sub-castes specialise in a particular set of tasks for a certain amount of time. Members of the sub-castes may be responsible for, among others things, brood care, foraging or nest-guarding. In some ant groups worker sub-castes are morphologically distinct and display, at the least, a clear size polymorphism (Buckingham, 1911; Detrain and Pasteels, 1992). In other highly eusocial taxa worker sub-castes are monomorphic and task specialisation is determined by age (Cameron, 1989). In primitively eusocial taxa, such as the paper wasp *Polistes*, female adult castes are behaviourally distinct but monomorphic and behaviourally plastic, meaning an adult worker can potentially become the dominant, reproducing queen at any time by replacing the current queen or founding a new colony (Sumner et al., 2006; Reeve et al., 2000).

These distinct morphological and behavioural castes, which exist among adult females of a eusocial colony, are based on alternative expression of the same genome. The plasticity of the behavioural castes in the primitively eusocial paper wasp, *Polistes canadensis*, was demonstrated by the existence of overlapping gene expression patterns along a continuum from newly emerged females, through intermediate workers to the dominant queens (Sumner et al., 2006). Most gene expression studies in this area have, however, concentrated on highly eusocial taxa. Large differences in gene expression have been recorded both between the morphologically distinct queens and workers (*Temnothorax longispinosus*: Feldmeyer et al. 2014; *Vespula squamosa*: Hoffman and Goodisman 2007; *Solenopsis invicta* & *S. richteri*: Ometto et al. 2011; *Apis mellifera*: Grozinger et al. 2007) and between monomorphic, behavioural worker sub-castes (*Temnothorax longispinosus*: Feldmeyer et al. 2014). The expression patterns of reproductive workers, that lay unfertilised eggs later in a colony cycle, become more ‘queen-like’ but they still remain more similar to non-reproductive workers than queens (Grozinger et al., 2007; Feldmeyer et al., 2014). Of the many genes found to be involved in caste differentiation *vitellogenin* has perhaps received most attention and has been shown to be differentially expressed among female castes of the honey bee and several ant species (Amdam et al., 2003; Feldmeyer et al., 2014; Corona et al., 2013; Morandin et al., 2014). Often in such studies a heavy focus has been placed on adult female castes, however, little work has been done to elucidate expression differences of males, but see Nipitwattanaphon *et al*. (2014). The haploid males are both morphologically and behaviourally distinct from their sisters and mother, but, although they differ in their ploidy level, they otherwise share the same genes as other colony members and are therefore also alternative expressions of the same genome.

Bumble bees represent an interesting taxon to study the phenomenon of eusociality as they possess both highly eusocial characteristics and more primitive features. For instance, whether a female will become a queen or a worker is irreversibly determined during development, as is the case for highly eusocial taxa. However, although a clear size dimorphism exists between queens and workers, generally both female adult castes are morphologically similar as in primitively eusocial species. Workers take on distinct tasks within a colony but the division of labour is more plastic than is the case for higher eusocial bees and is generally not temporally fixed (Cameron, 1989). Furthermore, towards the end of the colony cycle the division of labour between workers and reproductive queens breaks down and queens and workers come into direct conflict over the parentage of males. At this stage some workers activate their ovaries and begin to lay eggs and in the process become highly aggressive towards each other and also the queen (Alaux et al., 2004; Bloch, 1999).

So far no broad-scale studies have been conducted, which focus on the expression patterns involved in caste determination within bumble bees, although two previous studies did present some caste specific genes (Pereboom et al., 2005; Colgan et al., 2011). Pereboom et al. (2005) investigated how and when females developed into queens or workers. They identified, using suppression subtractive hybridisation, 12 genes whose expression differed in the comparisons: (1) worker and queen 1st instar larvae; (2) worker and queen 4th instar larvae; (3) adult queens and workers; (4) reproductive and non-reproductive workers. Colgan et al. (2011), within their analysis of the bumble bee transcriptome, found a high number of transcripts (2,185) that differ in their expression between adult castes, genders and developmental stages but considered their results as preliminary due to a lack of replication (1 larva, 1 pupa, 2 adult workers, 1 adult male and 1 virgin queen).

Here, using RNA-seq, we investigate genes involved in caste determination within the buff-tailed bumble bee, *Bombus terrestris*. We compare expression patterns of reproductive workers with those of non-reproductive workers and queens to isolate genes which are important for the acquisition of fertility as well as genes which may control behaviour differences compared to non-reproductive workers. Because of the flexible, plastic nature of bumble bee worker sub-castes (Cameron, 1989), reproductive workers are capable of becoming more ‘queen-like’ not only in their fertility but also in their behaviour. We therefore test the hypothesis that there is a greater similarity in gene expression patterns between queens and reproductive workers compared to those found in less plastic highly eusocial species.

Furthermore, we explore genes that control the specific behaviour and morphology of males. We investigate the question when, during the ontogeny of a male bumble bee, is the difference in gene expression to workers the greatest? Is the male gene expression pattern more distinct during larval development when the gonads and imaginal discs are generated? Are more genes involved in the development of the adult morphology during the pupal phase? Or does indeed the development and control of distinct behaviours among adults require the most distinct gene expression pattern? To address these questions we compare gene expression patterns of males and workers both within larvae and pupae. In adults, we analyse differences in expression patterns between males, queens, reproductive workers and non-reproductive workers.

## Materials & Methods

### Colonies

Six young, commercially available *B. terrestris audax* colonies were obtained from Agralan Ltd. Initially the colonies consisted of a mother queen and 8 to 20 workers. All colonies were kept in wooden nest boxes with the inner dimensions of 24 × 16 × 13.5 cm. The bees were supplied with pollen (mixed polifloral pollen, www.naturallygreen.co.uk) and a sugar solution (BIOGLUC^®^, Biobest) *ab libitum*. All colonies were kept in identical conditions within the same room at 26^°^C and 60% humidity in constant darkness.

### Sampling

We aimed to collect samples from 11 different combinations of caste and developmental stage, each from 3 independent colonies. Within larvae and pupae these were workers, males and queens, while in adults we intended to collect males, reproductive workers, non-reproductive workers, mother queens and virgin queens.

Sampling was carried out under red light conditions. The gender of adults was determined by counting antennal segments (males: 13; females: 12) and checking for the presence or absence of a sting (Prys-Jones and Corbet, 1987), while queens were identified via their superior mass (adult workers ranged from 26 to 325 mg, male adults from 127 to 347 mg and adult queens from 616 to 1,191 mg). In order to identify reproductive adult workers, samples from each colony were anesthetised by cooling for approximately 10 minutes, and their abdomens were dissected to observe ovary development. In order to avoid loss of RNA, dissections lasted only a few seconds and samples were immediately snap frozen in liquid nitrogen. Workers with developed ovaries were labeled ‘reproductive’. The workers, in which ovaries were not visible, were categorised to be of ‘undetermined reproductive status’, because of the potential time-lag between the expression of reproductive genes and subsequent changes in ovary morphology.

For the sampling of workers, queens and males during larval and pupal stages the following protocol was followed. The colonies were photographed at regular intervals of one to two days to monitor the emergence of new batches and their development. With the term ‘batch’ we refer to a single cohort of offspring laid together. At intervals of at least three days larvae and pupae were sampled from each batch while ensuring at least half of each batch was allowed to develop to adulthood. We collected larvae from each of the four larva instar stages based on their weight according to Cnaani et al. (1997) and assuming male instar masses were similar to worker instars. Pupae were collected both shortly after pupation (pre-pupae) and later in pupal development when appendages were developed.

Importantly gender and caste of all sampled larvae and pupae were confirmed by isolating batches after pupation and sexing all emerging adults. Only if 100% of the unsampled adults emerging from a batch belonged to the same gender and caste would the samples from that batch be considered for analysis. All samples were collected between the hours of 9am and 5pm as soon as they became available. They were immediately weighed, snap-frozen in liquid nitrogen and then stored at -80^°^C.

Worker larvae and pupae were obtained from batches laid and reared in young colonies in the presence of the queen. After sufficient worker batches were available the mother queen was removed from each colony for sampling. All batches laid in the presence of the queen but hatched shortly before or after the removal of the queen were considered potential queen batches (Pereboom et al., 2005). Any batches which were laid after queen removal were considered male batches. Additional male larvae and pupae were reared by isolating two to three groups of five workers from each colony in separate, small Perspex boxes containing pollen, sugar water and cat litter. The majority of male larvae and pupae (mean 67.2% ± 8.4% SEM) and all male adults were sampled from the main colonies.

As samples of the first larval stage were not obtained for workers from three separate colonies, L1 samples were excluded from all libraries. Adult virgin queens were only obtained from one colony, and the batches from which they emerged also produced adult workers. Therefore, larvae and pupae were only confirmed as queens if (1) they were sampled from batches from which adult queens emerged, and (2) if they exceeded 500mg (no sampled male or worker larva, pupa or adult exceeded 420 mg).

### RNA extractions

Whole bodies were used for sampling for two reasons. First, we had no prior assumptions regarding the tissues, within which genes would be differentially expressed between castes, genders and developmental stages. Second, in order to detect as many differentially expressed genes as possible across all comparisons. All samples were homogenised directly from −80^°^C. This was done within the Eppendorf tube with a plastic pestle for most larvae and in a ceramic mortar and pestle for large larvae, all pupae, and all adults. The mortar was filled with liquid nitrogen to keep the samples frozen during homogenisation. This was not necessary for the homogenisations which took place in Eppendorf tubes as the process was completed quickly. Total RNA was extracted from all samples using a GenElute Mammalian Total RNA Miniprep kit (Sigma-Aldrich) following the manufacturers’ protocol. The quality and concentration of RNA were estimated with an Agilent 2100 Bioanalyzer.

### RNA library construction

A total of 27 RNA libraries were constructed that covered all 11 combinations of caste and developmental stage from 1 or 3 colonies (table 1). Based on the concentrations estimated with the Bioanalyzer the larval libraries were prepared so as to contain equal quantities of RNA from each of the three larval stages 2 - 4 and equal quantities per individual within each larval stage. The same was also true for pre-pupae and pupae within the pupal libraries.

**Table 1:** The 27 RNA libraries and the number of pooled individuals contained in each.

| Caste | Developmental stage | Colonies |  |  |  |
| --- | --- | --- | --- | --- | --- |
|  |  | 7 | 8 | 9 | 11 |
| Worker | Larva (L2-L4) | 18 | 13 | 18 |  |
|  | Pupa | 6 | 4 | 5 |  |
|  | Reproductive adult | 1 | 1 | 1 |  |
|  | Undetermined reproductive adult |  | 1 | 1 | 1 |
| Queen | Larva (L4) | 1 |  |  |  |
|  | Pupa | 3 |  |  |  |
|  | Mother queen | 1 | 1 | 1 |  |
|  | Virgin queen | 1 |  |  |  |
| Male | Larva (L2-L4) | 16 |  | 9 | 7 |
|  | Pupa | 4 |  | 9 | 3 |
|  | Adult | 1 |  | 1 | 1 |

### Sequencing & assembly

The 27 libraries were sequenced on three lanes of an Illumina HiSeq 2500 system in rapid mode at the Edinburgh Genomics facility of the University of Edinburgh. After quality control and raw read processing, the reads were mapped to the *B. terrestris* transcriptome, BT_transcriptome_v2 (Colgan et al., 2011), using bwa_0.6.1. Only reads which mapped uniquely were considered for further analysis. Counts per transcript were subsequently calculated for each library using custom scripts.

### Differential expression analysis

The Blast2GO java program (Conesa et al., 2005) was used to annotate the transcriptome with gene descriptions and Gene Ontology (GO) terms (blastx against the nr database with e < 0.001). Differential expression analyses were carried out with the DESeq package (1.16.0; Anders and Huber 2010) in R (3.1.1; Team 2012).

A neighbor-joining tree was created based on expression differences between each of the 27 libraries. The distance matrix for the tree was calculated with the DESeq package and contained euclidean distances between each library based on variance stabilization transformed counts. The tree was created with Phylip (3.695, Felsenstein 2005). A principle components analysis was performed on all adult libraries within the DESeq package on variance stabilisation transformed data. Euler diagrams were created with the R package venneuler (Wilkinson and Urbanek, 2011).

Transcripts with a total of 50 reads or less across all 27 libraries were removed before performing the differential expression (DE) analyses. All remaining transcripts were tested for differential expression in each comparison. For each DE analysis, standard comparisons were performed between two conditions on normalized count data and with dispersion accounted for. Only transcripts with a Benjamini-Hochberg corrected p value (FDR) < 0.05 were considered as significantly, differentially expressed. For comparisons between castes within developmental stages, colonies were considered as replicates. No comparisons were made against queen pupae, queen larvae or adult virgin queens, as in each case only one replicate existed. These libraries were, however, included in comparisons of expression between developmental stages.

Gene function enrichment analyses (Fisher exact test) were carried out on DE transcripts with the R package topGO (2.16.0; Alexa and Rahnenfuhrer 2010). Enriched GO terms (FDR < 0.01) were subsequently summarised to meaningful clusters using Revigo (Supek et al., 2011). This method reduces redundancy of GO terms.

## Results

### Assembly

A total of 469.3 million 50 base pair, single-end reads were generated, ranging from 13.9 to 23.7 million reads per library. The reads mapped to the *Bombus terrestris* transcriptome at an average of 85.27% (75.25% to 92.47%) per library. The transcripts ranged in length from 101 to 26,110 bases (mean: 1,102; median: 721; fig. 1). Average read depth across the 27 libraries ranged from 0 to 47,420 (mean: 181; median: 16; fig. 1). All transcripts, to which a total of 50 or less reads (10,089, 27.8%) had been mapped across all libraries, were removed, leaving 26,265 (72.2%) transcripts for the differential expression analyses. The normalized counts per transcript ranged from 0 to 391,971 (median 34.41, mean 247.81) per library.

**Figure 1:**
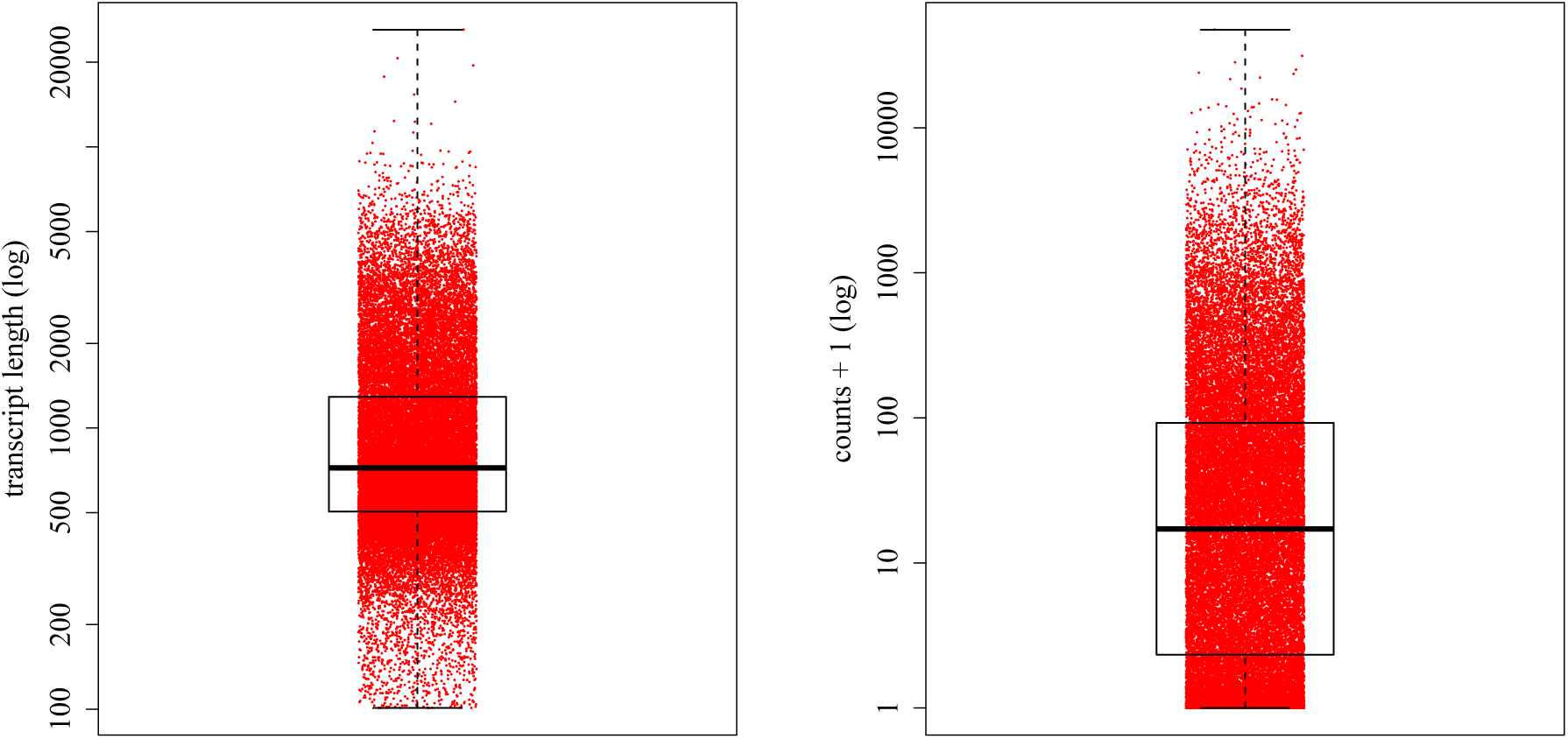
Range of transcript length in the transcriptome used for the assembly (left) and the average read depth per transcript across 27 libraries as raw read counts plus 1 (right). A log scale is used in both plots.

### Overview of gene expression patterns

All replicates, i.e. libraries from the same caste and developmental stage but from different colonies, showed low variation in their gene expression patterns and thus grouped together well in a neighbour-joining tree (fig. 2). The main clusters in the tree were formed by developmental stage (larvae, pupae and adults) rather than by caste. A differentiation in expression pattern between genders becomes more apparent in adults, where males form a distinct cluster. Within female adult castes a further clustering seems to have occurred. All reproductive workers and mother queens clustered together, and two of the workers with undetermined reproductive status (W*_Au_*8 and W*_Au_*11) formed a separate branch, while the adult virgin queen remained more distant to all other female adult groups. The adult worker with undetermined reproductive status from colony 9 (W*_Au_*9), on the other hand, grouped together with reproductive workers and mother queens. A principal component analysis (PCA) performed on all adult libraries indicated that W*_Au_*9 was indeed reproductive although ovaries had not been visible (fig. 3). In the analysis W*_Au_*9 clusters strongly with all reproductive workers and mother queens. W*_Au_*8 and W*_Au_*11 form a distinct group, well separated from the reproductive workers and mother queens. This clustering pattern could mean that W*_Au_*9 was reproductive and ovaries had not yet been visible in dissection. W*_Au_*8 and W*_Au_*11 were most likely non-reproductive.

**Figure 2:**
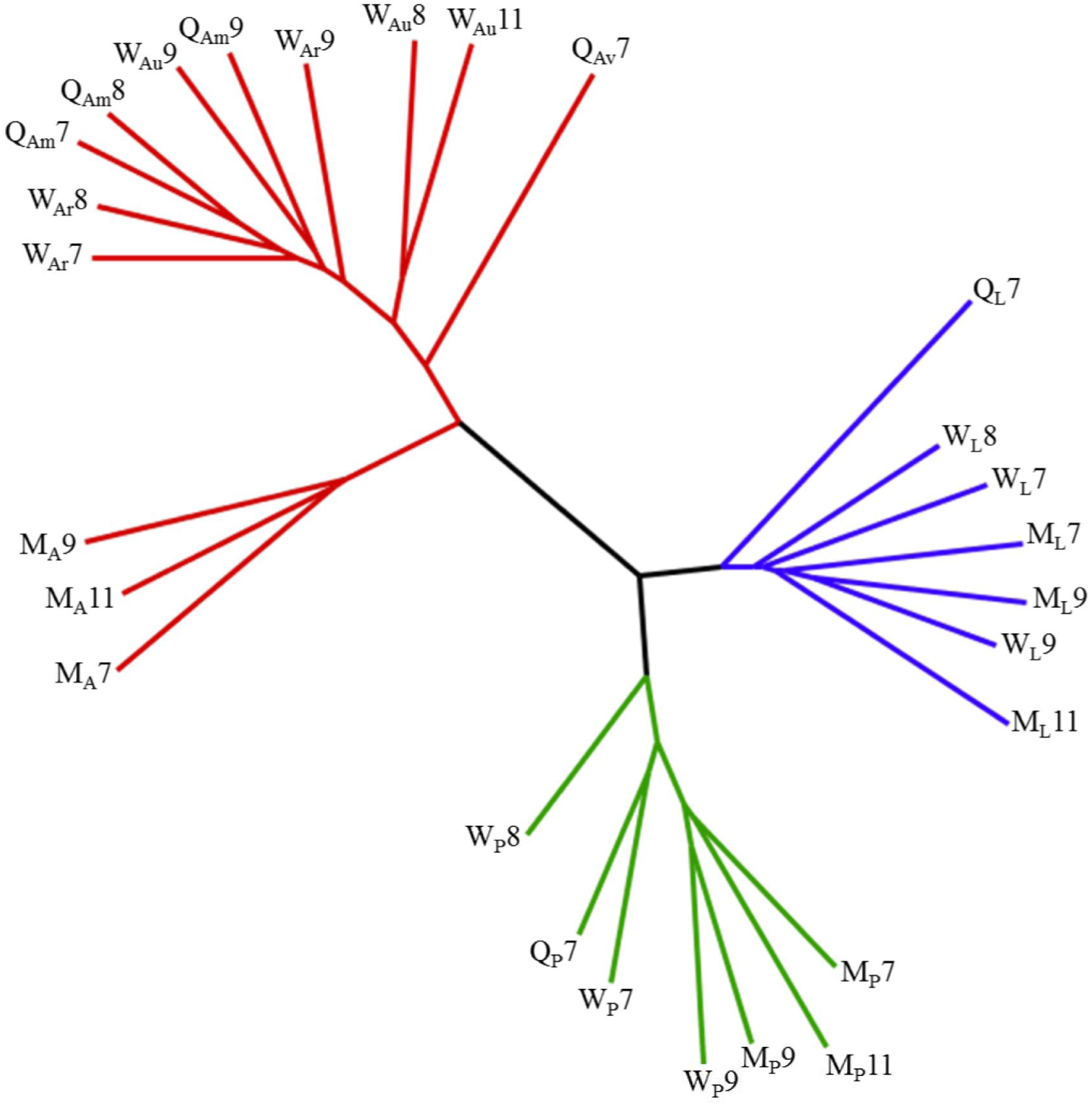
Neighbour-joining tree representing relationships between colonies, developmental stages, genders and castes based on expression pattern. Distances are euclidean and based on variance stabilization transformed counts. Numbers represent colonies; M = male; Q = queen; W = worker; _L_ = larva; _P_ = pupa; _A_ = adult; _r_ = reproductive; _u_ = undetermined reproductive status; _m_ = mother; _v_ = virgin.

**Figure 3:**
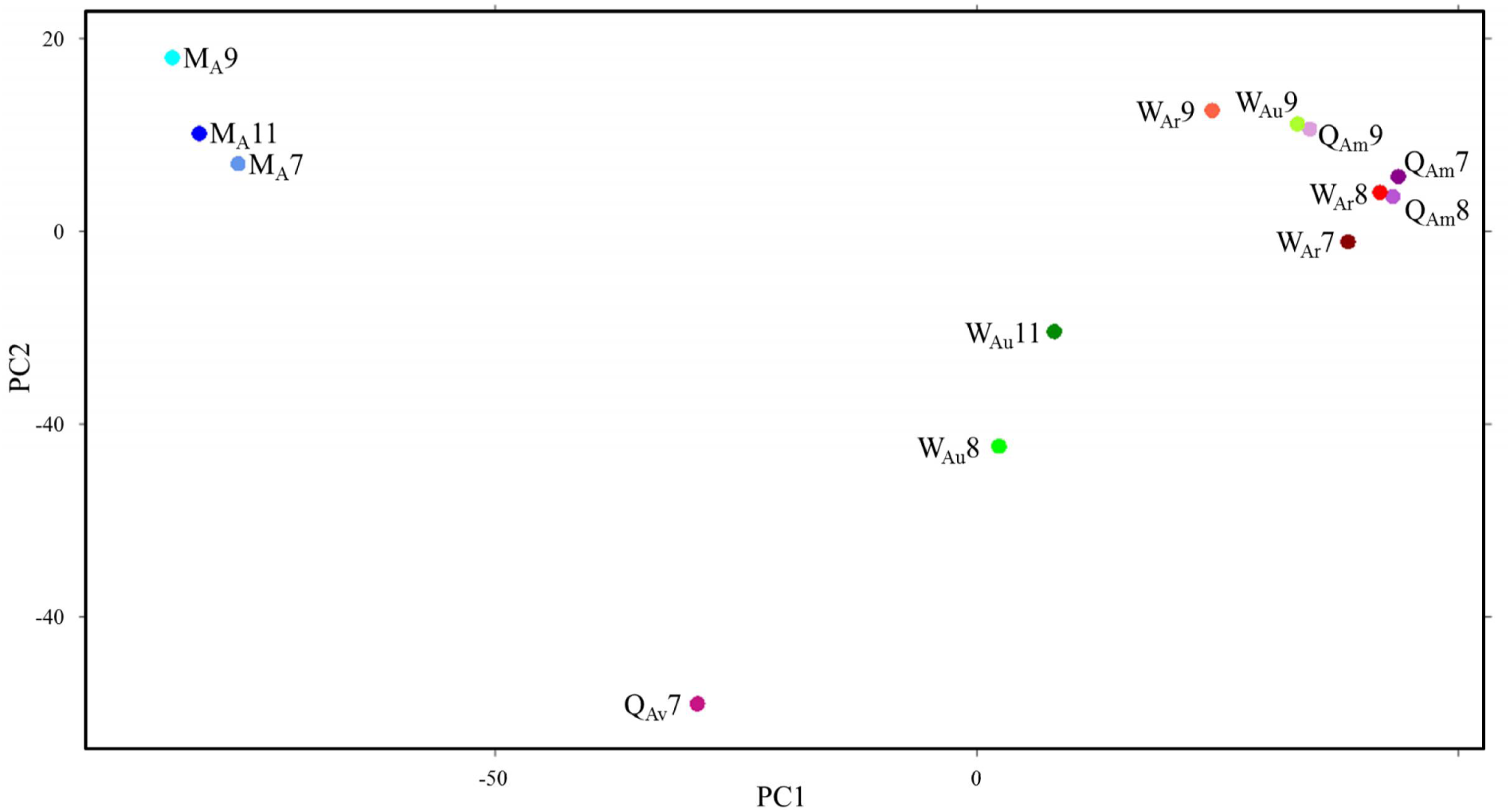
A principle components analysis of expression patterns among adult castes. The first two components explain 75.8% of variance. Distances are euclidean and based on variance stabilization transformed counts. Numbers represent colonies; M = male; Q = queen; W = worker; _L_ = larva; _P_ = pupa; _A_ = adult; _r_ = reproductive; _u_ = undetermined reproductive status; _m_ = mother; _v_ = virgin.

These conclusions were further supported by an over-representation of *Apis mellifera* reproductive genes (Cardoen et al., 2011) within W*_Au_*9 but not in W*_Au_*8 or W*_Au_*11. The expression of the 299 genes, which were over-expressed in reproductive honey bee workers compared to non-reproductive workers, was significantly lower in W*_Au_*8 (median 122.8; mean 363.2) and W*_Au_*11 (median 117.2; mean 389.4) than all three of our reproductive workers (W*_Ar_* 7: median 194.0, mean 638.9; W*_Ar_* 8: median 212.3, mean 627.6; W*_Ar_*9: median 138.9, mean 457.0). These differences were significant compared to W*_Ar_*7 (compared to W*_Au_*8: p = 0.0045; compared to W*_Au_*11: p = 0.0083) and W*_Ar_*8 (compared to W*_Au_*8: p = 0.0080; compared to W*_Au_*11: p = 0.0018; Mann-Whitney U test; fig. 4). Expression of the Cardoen reproductive genes was significantly higher in W*_Au_*9 (median: 196.5; mean: 542.5) than in both W*_Au_*8 and W*_Au_*11 (p = 0.0238 & 0.0376 respectively; Mann-Whitney U test; fig. 4). For these reasons W*_Au_*9 was considered reproductive and W*_Au_*8 and W*_Au_*11 were classed as non-reproductive for all further analyses.

**Figure 4:**
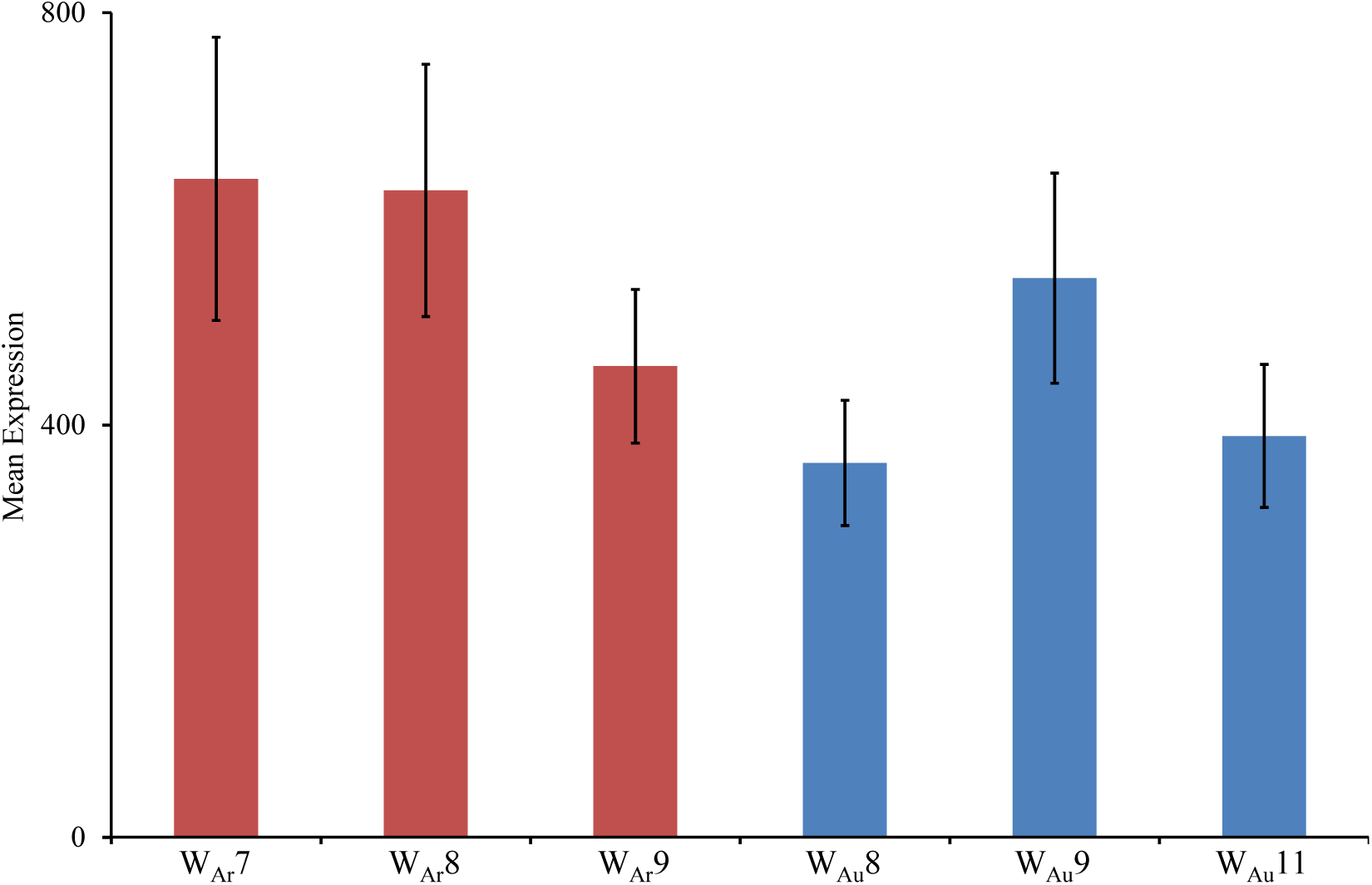
Mean expression level of honey bee reproductive genes identified in the study by Cardoen *et al*. (2011) in bumble workers of the present study. N = 299; error bars are standard error of the mean; W = worker; _A_ = adult; _r_ = reproductive; _u_ = undetermined reproductive status; numbers = colony.

The patterns shown in the neighbour-joining tree (fig. 2) and PCA of adult castes (fig. 3) were reflected in the number of DE transcripts found between developmental stages and castes. From 6,289 to 7,483 (mean 7,019) transcripts were differentially expressed between developmental stages. Only 71 and 162 DE transcripts were found between males and workers within larvae and pupae respectively, while a mean of 4,114 DE transcripts were found within adult comparisons ranging from 111 between reproductive workers and mother queens to 8,706 between adult males and mother queens (fig. 5).

**Figure 5:**
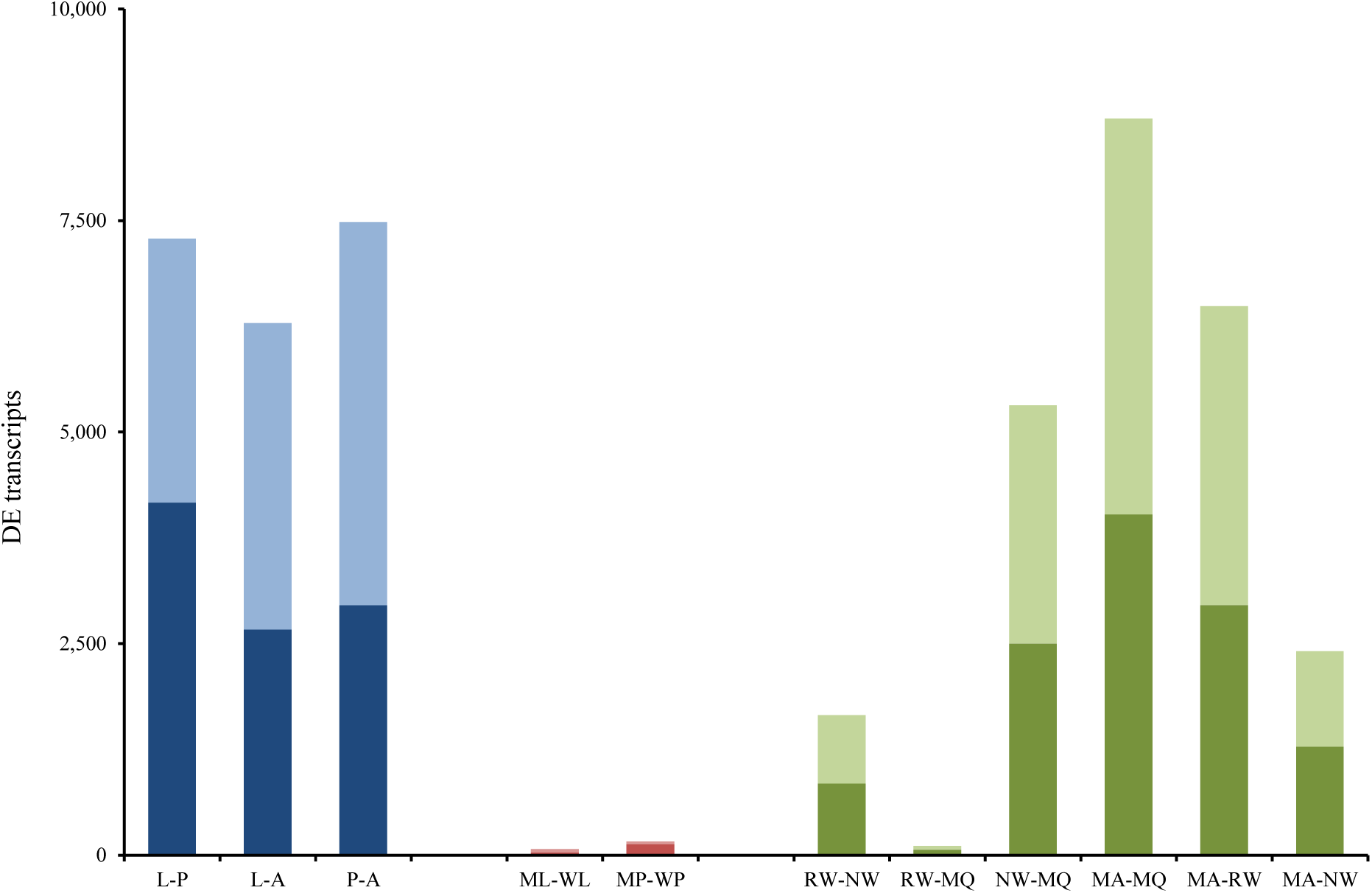
Differentially expressed transcripts within and between developmental stages. Darker colours: up-represented in first named caste; lighter colours: up-regulated in second named caste. M = male; W = worker; MQ = mother queen; L = larva; P = pupa; A = adult; R/N = reproductive/non-reproductive.

For some analyses of differential expression, colonies were not uniformly distributed, e.g. adult males (colonies 7, 9 & 11) versus non-reproductive workers (colonies 8 & 11). For these, ANCOVAs were performed to test for significant colony effects (see supplementary material). In only one out of 9 cases was a significant main effect of colony found. There were 5 cases where a significant interaction between colony and expression was found. In each of these cases a separate significant effect was still found for the important group difference: caste, developmental stage or gender. This means that any significant effects listed in the comparisons below can be attributed to differences in caste, gender or developmental stage rather than a colony effect.

### Developmental stages

A total of 12,218 DE transcripts were recorded in the three comparisons between larvae, pupae and adults (fig. 6). As already suggested by the neighbour-joining tree (fig. 2), adults differed most greatly from the other two developmental stages, confirmed by 3,237 transcripts which were differentially expressed compared to both pupae and larvae. A Gene Ontology (GO) term enrichment analysis showed that a heightened cellular metabolism distinguishes larvae from pupae and adults. The three main clusters of significantly over-represented GO terms (Fisher’s exact test, FDR <0.01) in a Revigo treemap were “translation”, “oxidative phosphorylation” and “ribosomal biogenesis” (fig. 7). Most over-represented GO terms among transcripts up-regulated in pupae either related to cell communication and movement, “signal transduction” and “cellular component organisation”, or the development of morphological features, “anatomical structure morphogenesis” (fig. 8). Most enriched adult GO terms belonged to the supercluster “G-protein coupled receptor signaling pathway” (fig. 9). This cluster included sub-clusters such as, “phototransduction”, “detection of stimulus” and “cell surface receptor signaling pathway”, highlighting the higher sensory capabilities and requirements of adults. 42.6% of larval, 58.7% of pupal and 48.3% of adult DE transcripts either received no significant blast hit or were linked to genes of unknown function. Detailed results of the Fisher tests can be found in the supplementary material.

**Figure 6:**
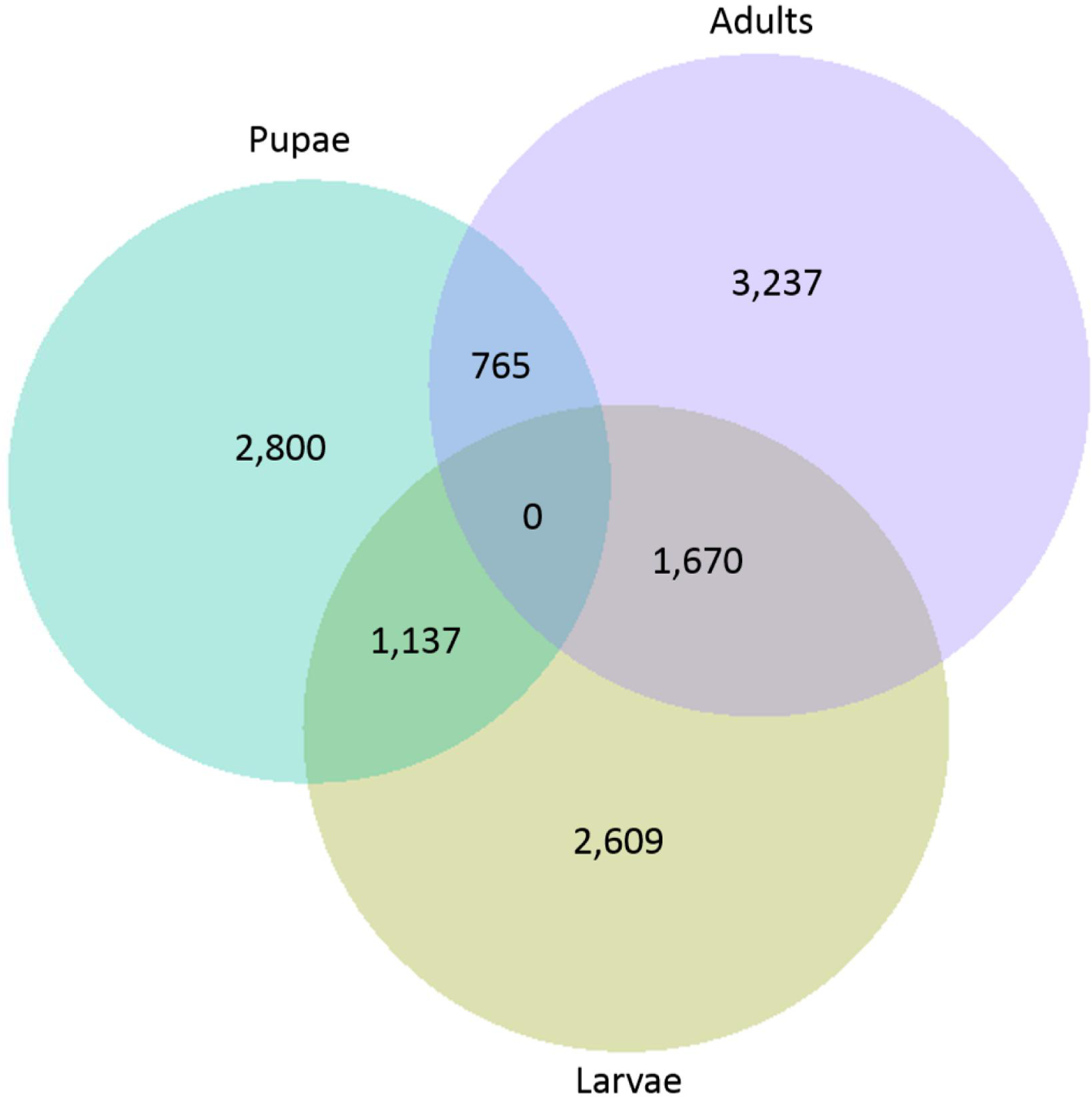
Number of transcripts which were differentially expressed between developmental stages.

**Figure 7:**
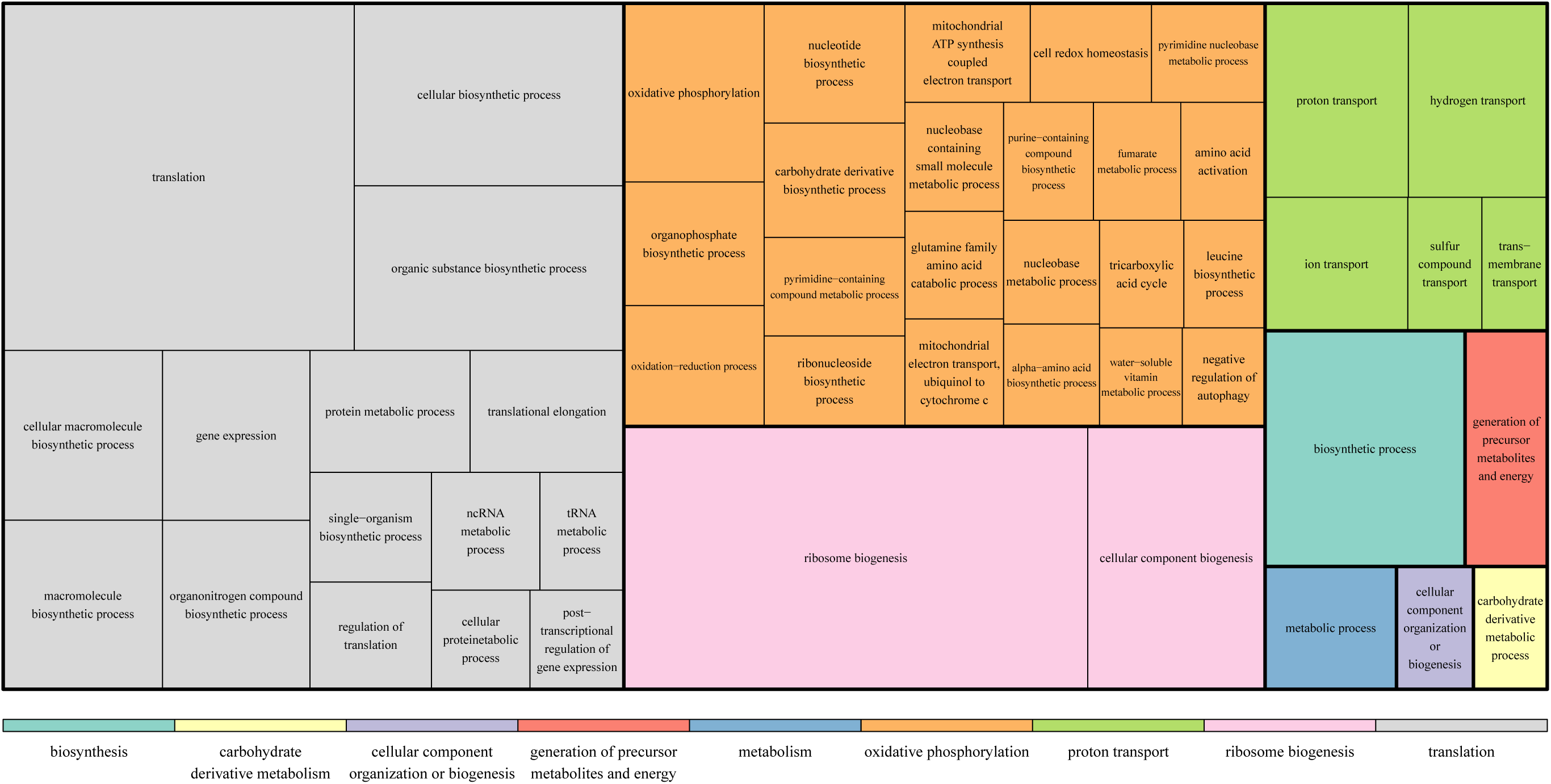
Most highly represented GO terms within larval DE genes (compared to pupae and adults).

**Figure 8:**
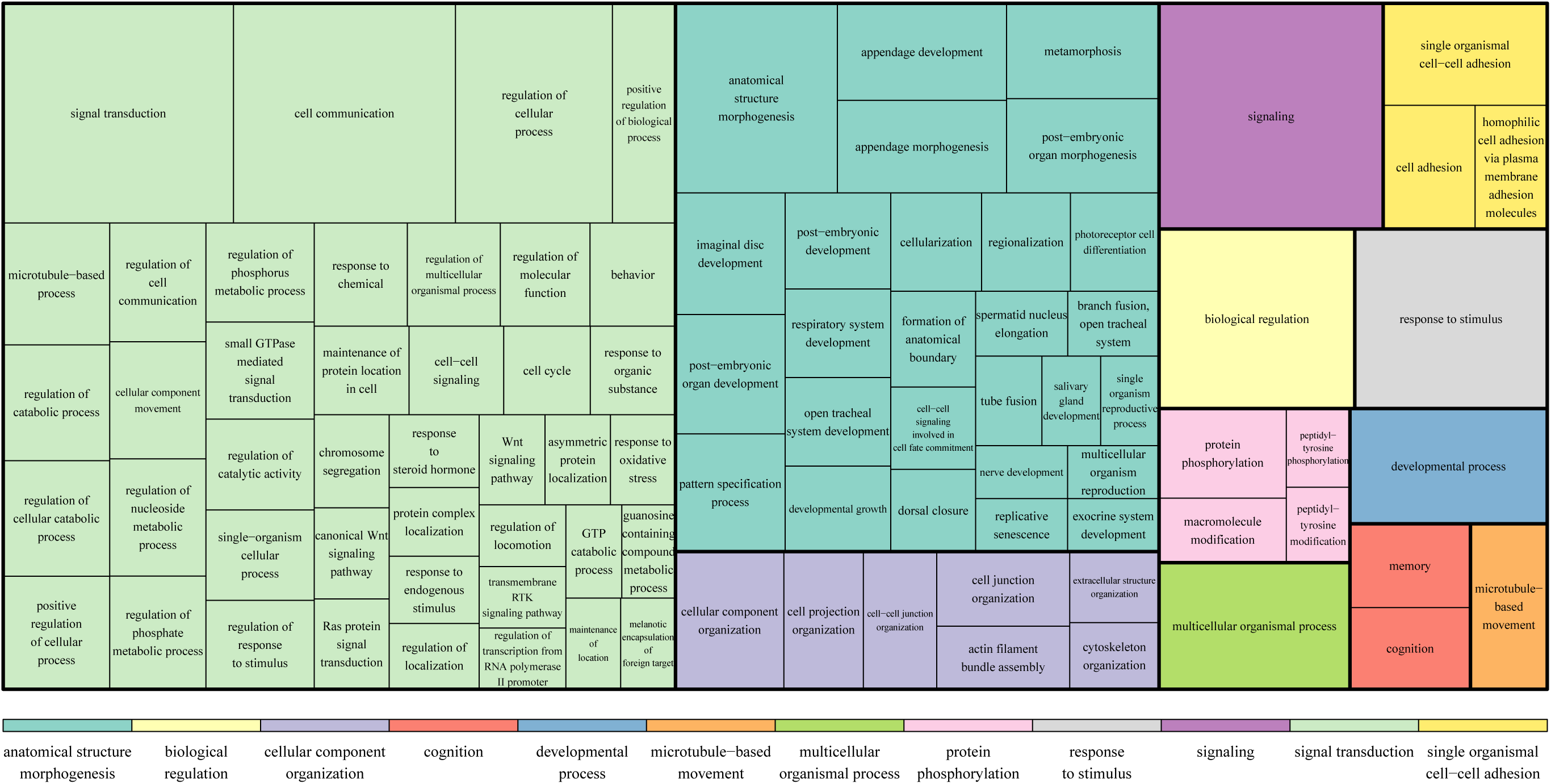
Most highly represented GO terms within pupal DE genes (compared to larvae and pupae).

**Figure 9:**
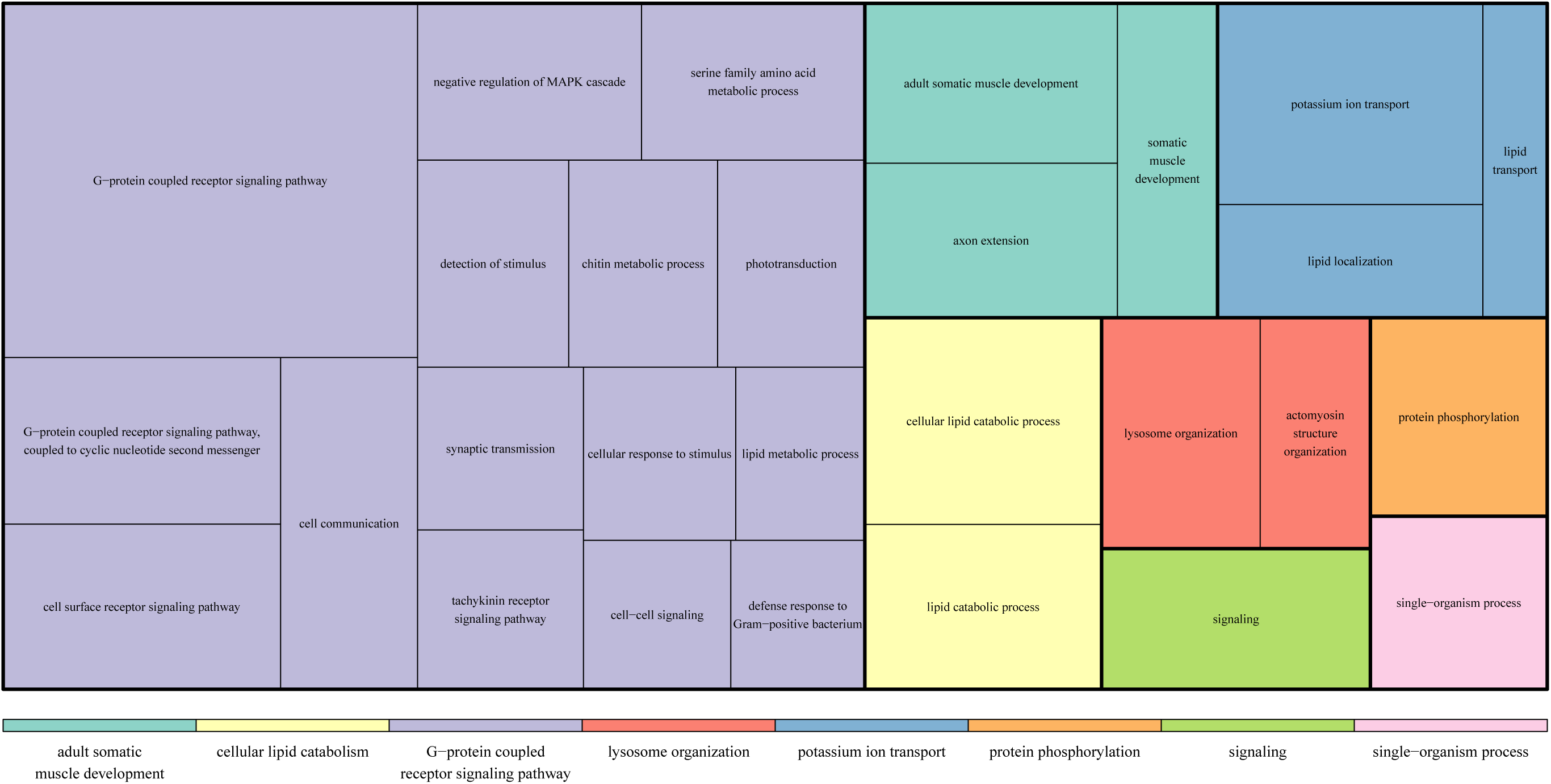
Most highly represented GO terms within adult DE genes (compared to larvae and pupae).

This test was repeated using lists of unique genes rather than transcript lists. All tendencies and the largest GO clusters remained unchanged. However, the number of significantly enriched GO terms was reduced. This was most likely due to the reduced number of genes in the test as a consequence of a high number of transcripts without an annotated gene match.

### Male versus worker larvae

Within larvae only a relatively small group of transcripts proved to be differentially expressed between males and workers (32 and 39 up-regulated transcripts respectively). Within the list of male larvae DE transcripts *nose resistant to fluoxetine protein 6-like*, *nrf-6*, appeared six times with a fold change (FC) ranging from 3.86 - 25.34 and expression of 48 to 4,576 mean normalised counts (mnc; supplementary tables). Nrf-6 is a transmembrane protein present in the intestine of various invertebrates (Choy and Thomas, 1999; Yao et al., 2014) and has been reported as up-regulated in the gut of *Ostrinia nubilalis* larvae (Lepidoptera) in response to a bacterial toxin (Yao et al., 2014). The presence of a further transcript within this list which encodes *cytochrome p450 6k1-like* (BTT39618_1; 2.07 FC; 1,682 mnc; supplementary tables) provides possible further evidence for an infection within the male larvae. Riddell et al. (2014) found in *B. terrestris* that the expression of 16 different cytochrome p450 transcripts was altered post infection.

*Takeout-like* (XP_003397291.1; transcript BTT15842_1) was also strongly up-regulated in male larvae compared to worker larvae (5.75 FC; 2,693 mnc; supplementary tables). A close homolog to this transcript (blastp: 68% identity, e-value 3e^−126^) has been characterized for *A. mellifera* (Hagai et al., 2007). Takeout (to) was reported to be involved in the regulation of maturation in worker honey bees. In that study only adult workers were investigated so that any gender or developmental effects are as yet unknown for Hymenoptera. However, the *to* gene family is known to be over-expressed in adult *Drosophila* males, affecting courtship behaviour (Dauwalder et al., 2002).

The majority of the worker larvae DE genes (24 out of 39; 8 of the top 10 in terms of FC) were either of unknown function or received no significant blast hits (supplementary tables). One *vitellogenin* transcript (BTT24408_1; 4.35 FC; 62 mnc; supplementary tables) was over-expressed in worker larvae compared to male larvae.

### Male versus worker pupae

Differentiation was somewhat greater between males and workers during the pupal phase compared to the larval phase. 128 transcripts were significantly up-regulated in male pupae and 34 in worker pupae. The pupal list contained a high number of uncharacterized transcripts: 84 (66%) male and 24 (71%) worker pupae transcripts (supplementary tables).

Six male DE transcripts coded for *tubulin* related genes (3 *α*-tubulin transcripts, 1 *β*-tubulin transcript and 2 tubulin-tyrosine ligases; 6.28 - 490.54 FC; 56 - 1,200 mnc; supplementary tables). The tubulin-tyrosine ligase is involved in the post-transcriptional modification of *α*-tubulin (Ersfeld et al., 1993), so it appears tubulin transcripts, especially *α*, may be important for male pupal development. The same *vitellogenin* transcript up-regulated in worker larvae (BTT24408_1) was also up-regulated in worker pupae compared to male pupae (5.83 FC; 29 mnc; supplementary tables).

### Fertility genes

Within the comparisons between adult castes (males, reproductive workers, non-reproductive workers and mother queens), reproductive workers and mother queens were most similar with only 111 DE transcripts (64 up-regulated in reproductive workers and 47 in mother queens; fig. 5). Non-reproductive workers, on the other hand, were distinct from both mother queens (2,499 up-regulated in non-reproductive workers, 2,817 in mother queens) and, to a lesser extent, reproductive workers (844 up-regulated in reproductive, 810 in non-reproductive workers). The majority (791, 93.7%) of the transcripts up-regulated in reproductive workers compared to non-reproductive workers were also up-regulated in mother queens compared to non-reproductive workers. As the common difference between non-reproductive and reproductive workers and between non-reproductive workers and mother queens is their fertility status, we have named these 791 transcripts ‘fertility genes’ (fig. 10).

**Figure 10:**
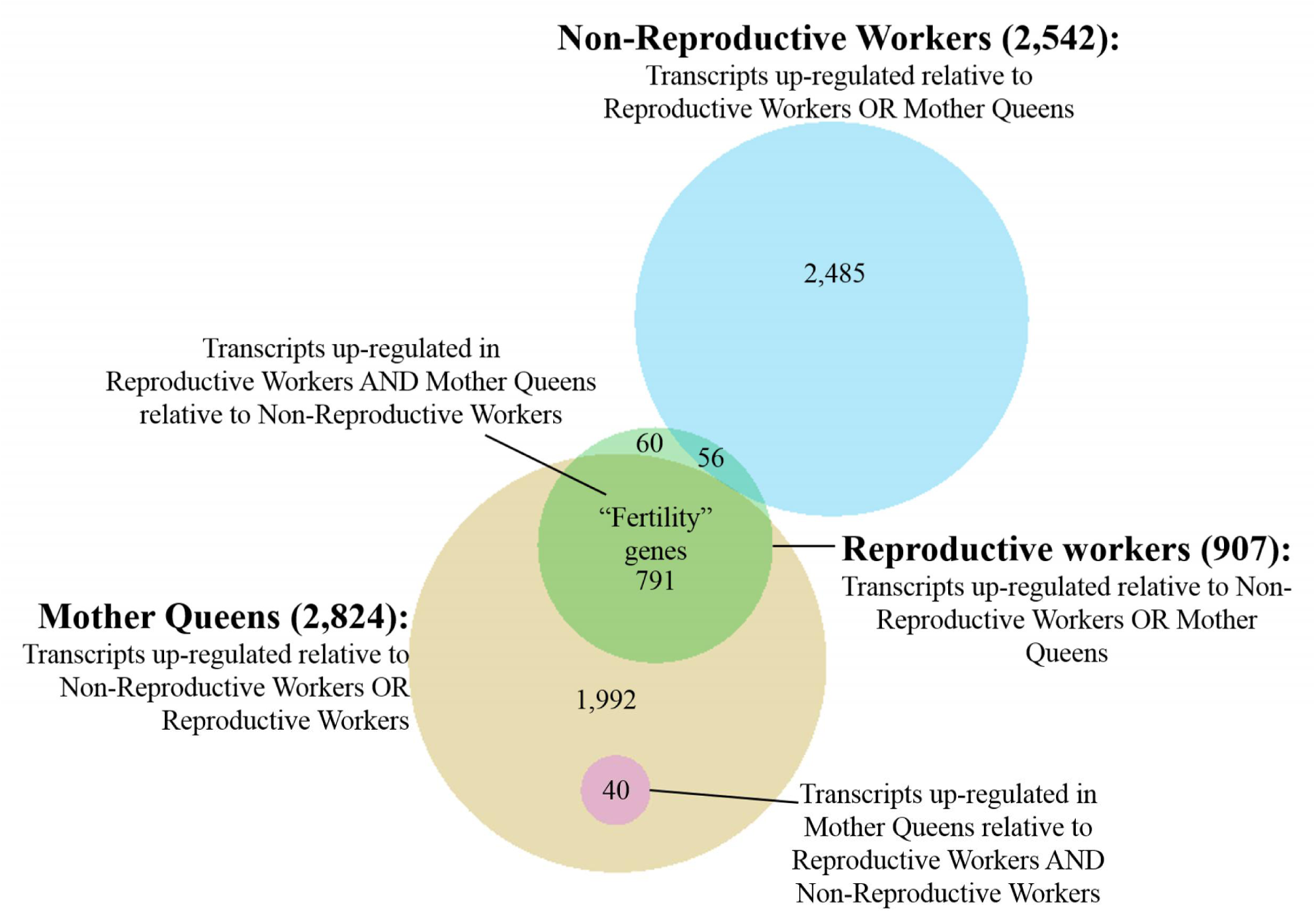
The number of transcripts which are differentially expressed between female adult castes. Overlapping areas indicate shared transcripts.

All differential expression values in this section are based on the comparison of reproductive and non-reproductive workers, although all transcripts were also up-regulated in mother queens versus non-reproductive workers. 267 (33.8%) of the fertility transcripts were of unknown function (1.78 - 336.18 FC; 18 - 39,927 mnc; supplementary tables). A large number of transcripts were involved in protein synthesis activity: a total of 54 up-regulated gene transcripts contained the labels “transcription”, “translation”, “RNA polymerase”, “ribosomal”, “ubiquitin”, “helicase” or “ribonucleoprotein” (1.85 - 11.68 FC; 68 - 29,132 mnc). Seven *tudor* transcripts, a gene known to be involved in the formation of female germ cells in *Drosophila* (Boswell and Mahowald, 1985), were significantly higher expressed in reproductive workers with a fold change ranging from 1.88 to 2.44 (144 - 491 mnc). 61 of the fertility transcripts (1.77 - 9.53 FC; 12 - 9,973 mnc) were direct homologs of genes up-regulated in honey bee reproductive workers in a similar comparison (Cardoen et al., 2011). These transcripts encoded genes with functions such as oocyte meiosis, oocyte axis specification, oogenesis and female gonad development (supplementary tables).

The list also contained two *vitellogenin* (4.95 & 6.03 FC; 4,103 & 111,595 mnc) and four vitellogenin receptor transcripts (1.94 - 3.39 FC; 222 - 11,577 mnc). The two *vitellogenin* transcripts had, on average across all libraries, a total expression level of 45,294 mnc, making up 69.4% of all *vitellogenin* transcripts on average per individual (97.7% in mother queens and 98.4% in reproductive workers; fig. 11a, b). The *vitellogenin* transcripts (BTT24408_1 and BTT40935_1) are closely related to the 1,772 amino acid vitellogenin genes ACQ91623 and ACU00433 of *B. ignitus* and *B. hypocrita* respectively (table 2). These genes correspond to the conventional *Vg1* gene described by Morandin *et al*. (2014; blastp: E = 0.0, Id = 33%). The four receptor transcripts corresponded to the two *B. terrestris* genes *vitellogenin receptor-like isoform 1* and *isoform 2* (XP_003402703 and XP_003402704).

**Figure 11:**
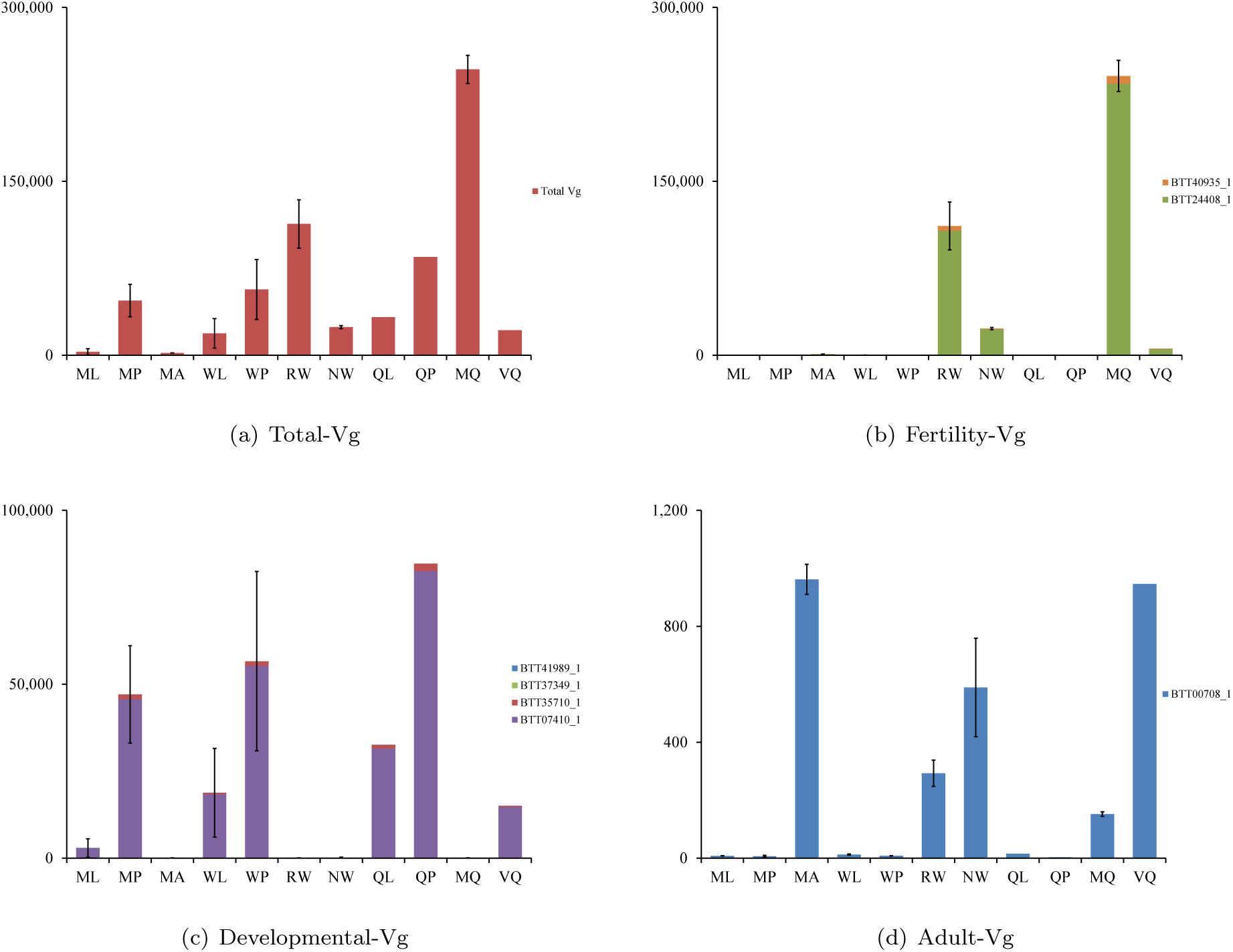
Vitellogenin expression levels within different castes and developmental stages. Expression is mean of normalized counts across replicates; error bars are standard error of the mean. (a) the summed expression level of 18 vitellogenin transcripts; (b) 2 vitellogenin transcripts up-regulated in reproductive workers and mother queens versus non-reproductive workers;(c) 4 vitellogenin transcripts up-regulated in larvae and pupae versus adults; (c) 1 transcript up-regulated in all adults compared to larvae and pupae but down-regulated in reproductive adults. M=male, W=worker, Q=queen, L=larvae, P=pupae, A=adult, NR/R=non-/reproductive, M=mother, V=virgin; All castes include samples from 3 colonies, except RW: 4; NRW: 2; QL, QP & VQ: 1.

**Table 2:** 7 vitellogenin transcripts and the castes in which they are up-regulated.

| Transcript | Caste specificity | Top blastx hit | e value | Protein length | Morandin *et al.* 2014 | Mean expression across 27 libraries |
| --- | --- | --- | --- | --- | --- | --- |
| BTT24408_1 | Reproductive female adults | ACQ91623 (*B. ignitus*) - vitellogenin | 0.0 | 1,772 | *Vg1* | 43,917 |
| BTT40935_1 |  | ACU00433 (*B. hypocrita*) - vitellogenin | 5e<sup>-62</sup> |  |  | 1,377 |
| BTT07410_1 | Larvae and pupae | XP_003400264 (*B. terrestris*) vitellogenin-6-like | 0.0 | 1,514 | *Vg-like-A* | 18,374 |
| BTT35710_1 |  |  | 1e<sup>-164</sup> |  |  | 477 |
| BTT37349_1 |  |  | 0.0 |  |  | 9 |
| BTT41989_1 |  |  | 3e<sup>-29</sup> |  |  | 10 |
| BTT00708_1 | Adult males and non-reproductive adult females | XP_003393940 (*B. terrestris*) vitellogenin-like | 0.0 | 319 | *Vg-like-C* | 250 |

*Vitellogenin* was, however, not restricted to female reproductive castes. The second highest expressed *vitellogenin* transcript across all libraries, BTT07410_1, constituted on average 28.1% of vitellogenin transcripts. This transcript together with three further transcripts (BTT35710_1, BTT41989_1 and BTT37349_1) is associated with the *B. terrestris* gene XP 003400264 (*vitellogenin-6-like*), which is 1,514 amino acids in length and corresponds with the *Vg-like-A* gene described by Morandin *et al*. (2014; blastp: E = 0.0, Id = 44%; table 2). These four transcripts appear to be involved in development and independent of gender as they were up-regulated in all larvae and pupae samples compared to adults irrespective of caste and gender (fig. 11c). One *vitellogenin* transcript (BTT00708_1) was significantly up-regulated in adults compared to pupae and larvae but was down-regulated by reproductive workers (significantly compared to male adults) and mother queens (significantly compared to non-reproductive workers and male adults; fig. 11d). This transcript is coded by the *B. terrestris* vitellogenin-like gene XP_003393940, which is much shorter than the two previously discussed *vitellogenin* genes (319 amino acids) and is similar to *Vg-2* of *Apis mellifera* (blastp: 66% identity, e-value 1e^−142^) and the *Vg-C-like* homolog described in Morandin *et al*. (2014; blastp: E = 1e^−134^, Id = 57%; table 2).

Seven *α-glucosidase* transcripts were differentially expressed within the fertility genes (6.93 - 9.14 FC; 16 - 35,260 mnc). An analysis of all 10 *α-glucosidase* transcripts within the *B. terrestris* transcriptome across all libraries showed raised expression levels for reproductive workers and mother queens compared to all other castes and developmental stages. Non-reproductive workers had the third highest levels of the 11 combinations of caste and developmental stage but *α-glucosidase* transcripts were 8 times more abundant in reproductive workers and mother queens (fig. 12). Four *glucose dehydrogenase* transcripts (BTT01220_1, BTT08099_1, BTT18258_1 & BTT20465_1), on the other hand, were down-regulated in mother queens and reproductive workers, although up-regulated in all adults compared to larvae and pupae (fig. 13). These transcripts all related to the *B. terrestris glucose dehydrogenase* gene XP_003395668.1.

**Figure 12:**
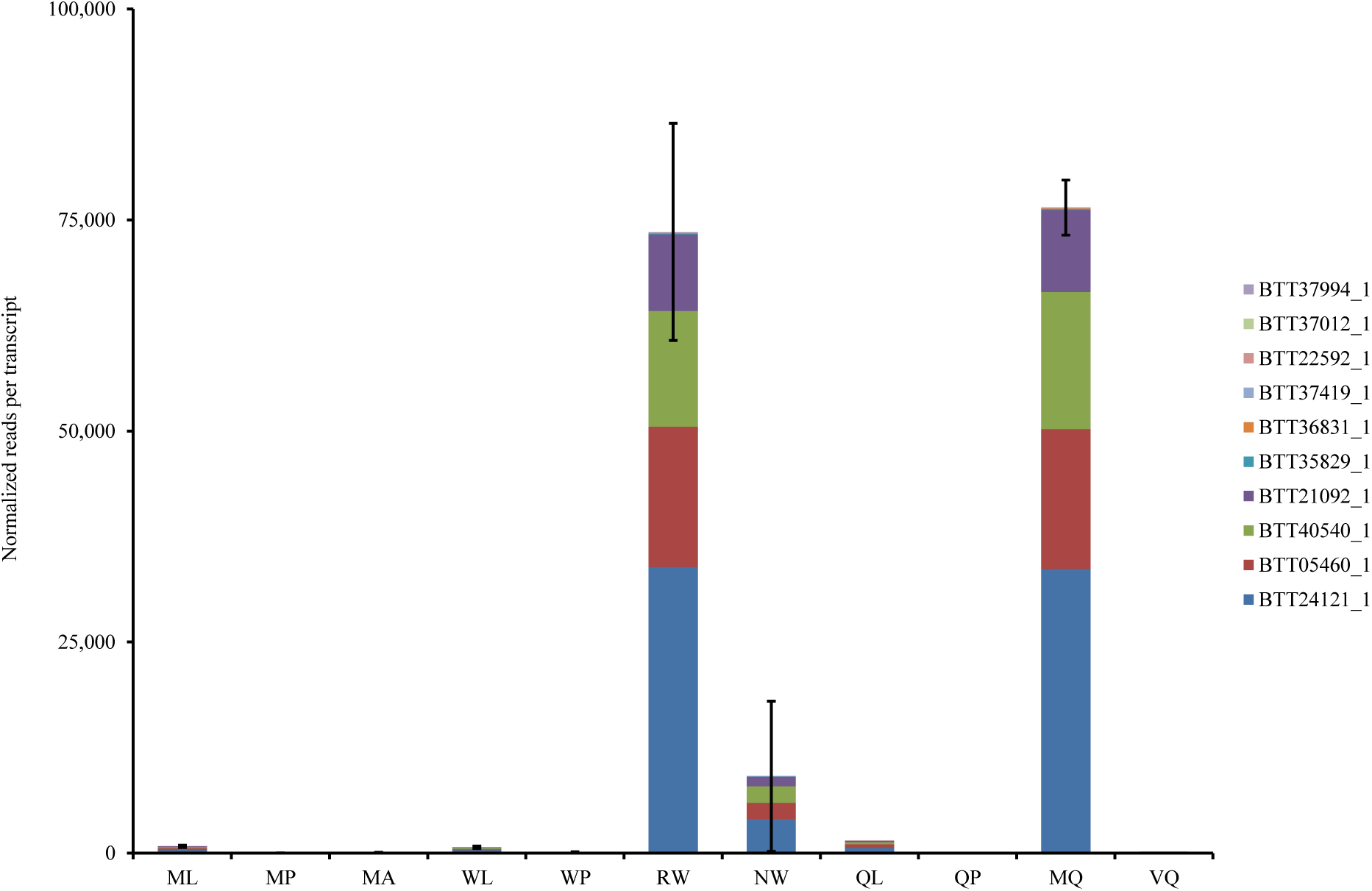
Expression levels of 10 *α*-glucosidase transcripts within 11 different castes or developmental stages. Expression is mean of normalized counts across replicates; error bars are standard error of the mean. M=male, W=worker, Q=queen, L=larvae, P=pupae, A=adult, NR/R=non-/reproductive, M=mother, V=virgin; All castes include samples from 3 colonies, except RW: 4; NRW: 2; QL, QP & VQ: 1.

**Figure 13:**
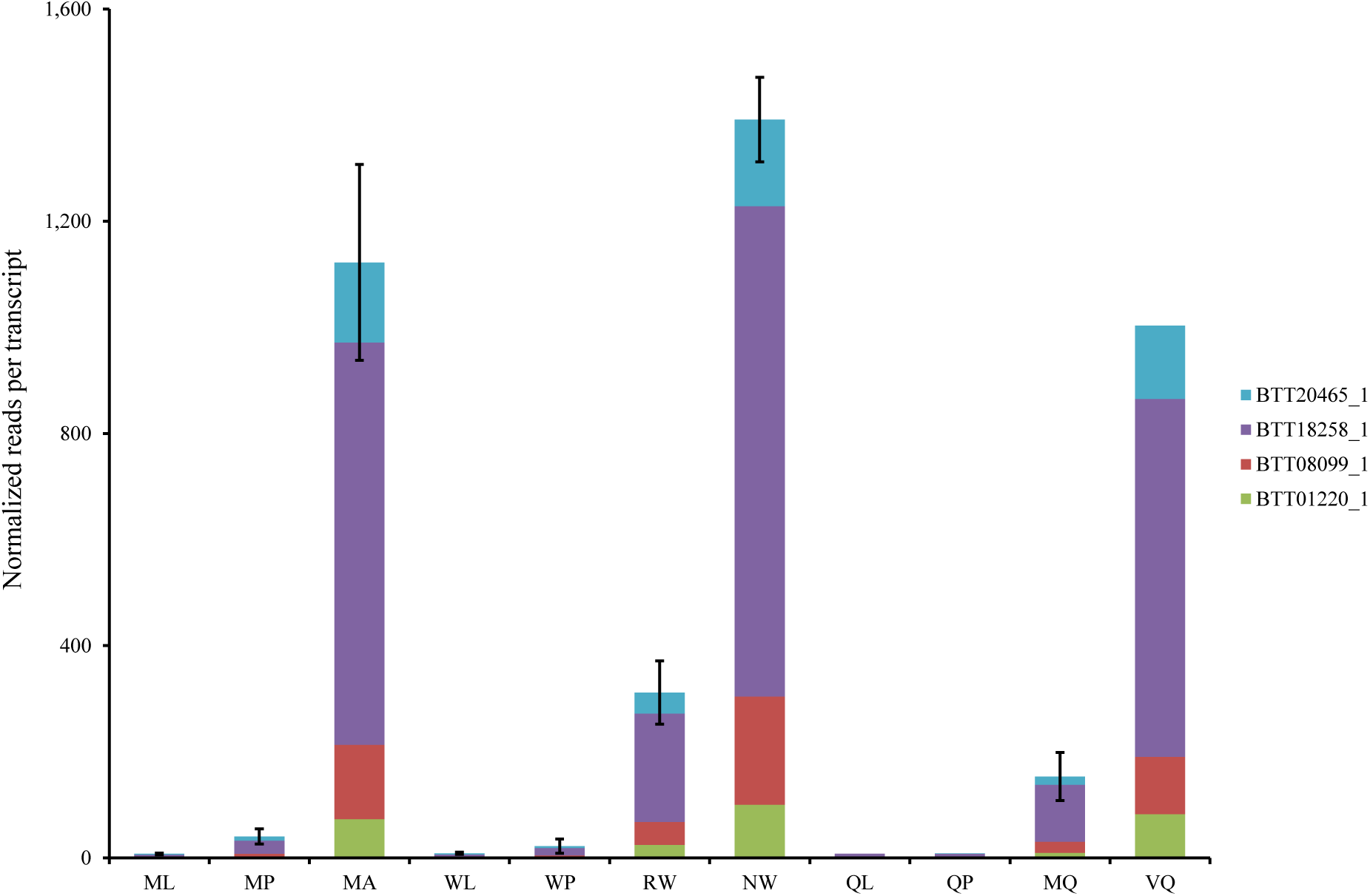
Expression levels of 4 glucose dehydrogenase transcripts within 11 different castes or developmental stages. Expression is mean of normalized counts across replicates; error bars are standard error of the mean. M=male, W=worker, Q=queen, L=larvae, P=pupae, A=adult, NR/R=non-/reproductive, M=mother, V=virgin; All castes include samples from 3 colonies, except RW: 4; NRW: 2; QL, QP & VQ: 1.

Interestingly, mean expression of the 10 *α-glucosidase* transcripts correlated significantly with mean expression of the two *vitellogenin* transcripts (BTT24408_1 and BTT40935_1), which were also up-regulated in the fertility genes (*ρ* = 0.7247; p = 1.91 × 10^−5^; Spearman’s rho). Similarly, mean expression of the four *glucose-dehydrogenase* transcripts, down-regulated in fertility genes, significantly correlated with the down-regulated *vitellogenin* transcript (BTT00708_1; *ρ* = 0.7888; p = 1.02 × 10^−5^; Spearman’s rho).

Two transcripts (BTT20241_1 & BTT33633_1; 67.16 & Inf FC; 5,528 & 37 mnc), which encode laccase-2-like, were up-regulated in reproductive versus non-reproductive workers but not in mother queens versus non-reproductive workers. Laccase 2 is a protein involved in the sclerotisation of extracellular structures in invertebrates (Arakane et al., 2005).

### Non-reproductive workers

For the majority (465 out of 810; 57.4%) of the transcripts up-regulated in non-reproductive workers compared to reproductive workers the function was unknown (supplementary tables). 19 of the non-reproductive worker genes were direct homologs of genes up-regulated in non-reproductive *A. mellifera* workers (Cardoen et al., 2011). Eight of those (1.77 - 3.74 FC; 72 - 653 mnc) had been attributed to the effect of the queen mandibular pheromone (QMP) in a previous study (Grozinger et al. 2003; supplementary tables).

### Adult queens

Transcripts, which were up-regulated in mother queens compared to both reproductive and non-reproductive workers, were considered ‘queen genes’ (fig. 10). The 40 queen transcripts ranged in fold change compared to reproductive workers from 1.68 to 8.87 (29 - 245,472 mnc; supplementary tables). Eleven of the transcripts (27.5%) were of unknown function. Most notable among the queen genes were 5 transcripts relating to serine protease inhibitors, SPI (2.92 - 8.87 FC; 2,145 - 10,596 mnc). These five SPIs were expressed together at a mean of 27,758 mnc ±1,247 SEM in mother queens compared to only 5,026 mnc in the virgin queen (fig. 14; supplementary tables). The second highest levels were found in non-reproductive workers (7,555 mnc ±1,527 SEM) followed by reproductive workers (6,435 mnc ±699 SEM).

**Figure 14:**
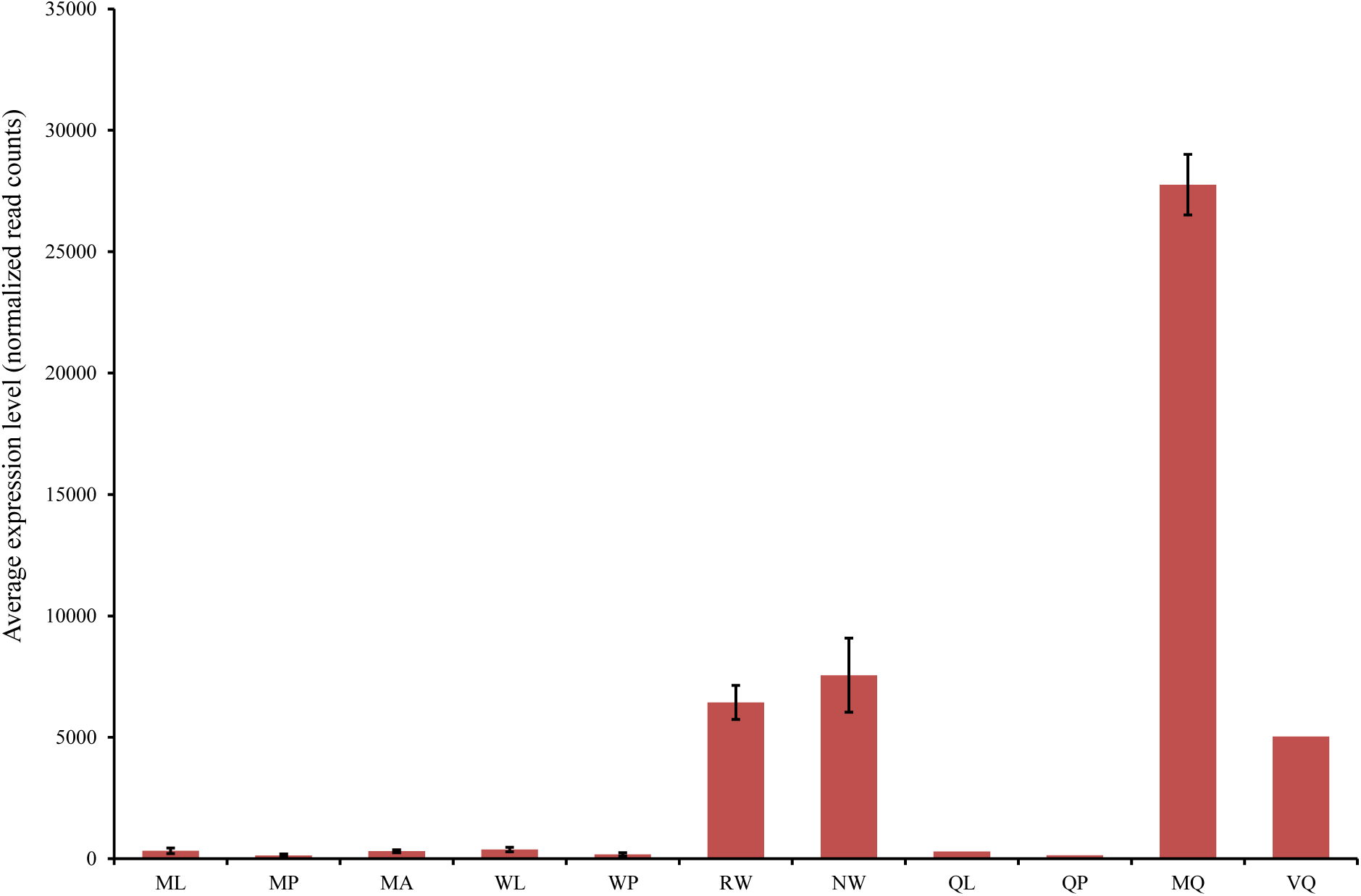
Expression levels of 6 serine protease inhibitors. Expression is mean of normalized counts across replicates; error bars are standard error of the mean. M=male, W=worker, Q=queen, L=larvae, P=pupae, A=adult, NR/R=non-/reproductive, M=mother, V=virgin; All castes include samples from 3 colonies, except RW: 4; NRW: 2; QL, QP & VQ: 1.

### Adult males

In males compared to non-reproductive workers 1,280 transcripts were up-regulated, of which 526 (41.1%) were of unknown function (supplementary tables). A high number of male transcripts (190), containing the tags “mitochond”, “cytochrome”, “pyruvate”, “NADH dehydrogenase” or “quinone”, were involved in the mitochondrial metabolism (1.85 - 41.66 FC; 8 - 62,872 mnc). 37 transcripts were involved in muscle development (*myosin*, *troponin*, *twitchin* and *titin*; 2.42 - 28.60 FC; 10 - 5,877mnc) and a further 16 in the fatty acid metabolism (1.94 - 202.24 FC; 6 - 3,935 mnc).

### Comparison with previous studies on Bombus terrestris

The top 10 transcripts up-regulated in larvae in the study carried out by Colgan et al. (2011) related to cuticle proteins, the storage protein hexamerin and the metabolic proteins carbonic anhydrase and cytochrome p450. In the present study 5 cuticle, 2 hexamerin (70c and 70b), 10 carbonic anhydrase and 12 cytochrome p450 related transcripts were also up-regulated in larvae compared to pupae and adults. The 10 transcripts listed in Colgan et al. could be linked to one GO term (GO:0042302: “structural constituent of cuticle”), which was also attributed to 17 of the larvae transcripts (up-regulated relative to pupae and adults). In a further study a cuticle protein and hexamerin were also present in larvae but absent in adults; pupae were not included in the analysis (Pereboom et al., 2005). The *vitellogenin* transcript BTT07410_1, which we found to be up-regulated in larvae and pupae, was also over-expressed in pupae in the Colgan et al. study (2011), however, not detected in larvae. All 7 of the GO terms, which were associated with the top 10 pupal genes in the Colgan et al. study (2011), were also present in our list of up-regulated pupae transcripts (p = 1.3 × 10^−4^, hypergeometric test).

In workers Colgan et al. (2011) found over-expressed genes associated with flight, defence and metabolism (*cytochrome p450*, *lipase* and *α-glucosidase*). In the present study flight muscles were also over-represented in non-reproductive workers and the metabolism genes *lipase*, *cytochrome p450* and *α-glucosidase* were more highly expressed in workers than in males. 20 of the 36 GO terms associated with the worker transcripts in the Colgan *et al*. study were also found in the transcripts up-regulated in non-reproductive workers relative to adult males in the current study (p = 5.8 × 10^−9^, hypergeometric test). The genes differentially expressed between adult female castes and sub-castes in the Pereboom et al. study (2005), *60-S ribosomal protein*, *chymotrypsin*, *cytochrome oxidase*, *peroxiredoxin*, *fatty acyl CoA-desaturase* and *ATP synthase beta subunit*, could not be confirmed with our data.

Colgan et al. (2011) found transcripts of the flight muscle gene *titin* to be over-represented in male adults, as well as several immunity genes. Many flight muscle proteins were also up-regulated in our study, however, we could not confirm the over-representation of immunity genes among the transcripts with known function. All 17 GO terms present in the top 10 male transcripts of the Colgan et al. study (2011) were also present in our list of male transcripts (up-regulated in adult males relative to non-reproductive workers; p = 2.3 × 10^−14^, hypergeometric test).

## Discussion

We compared gene expression patterns both between developmental stages and between castes within each developmental stage for the buff-tailed bumble bee *Bombus terrestris*. The number of differentially expressed transcripts ranged from 71 between male and worker larvae to 8,706 between adult males and mother queens. We found gene expression patterns to differ more between developmental stages than between caste or gender. Genes up-regulated in larvae were associated with a high cellular metabolism, whereas in pupae over-expressed genes were associated with cell communication and the development of morphological features. Most of the over-represented GO terms in adults were related to the G-protein coupled receptor signaling pathway. G-proteins are cell-surface receptors, which respond to extra-cellular stimulants with an intracellular signal cascade (Dohlman, 2002; Strader et al., 1994).

The number of genes differentially expressed became progressively larger through the three developmental stages as each caste became more distinct. These findings suggest a comparatively low number of genes are required to create distinct morphological castes compared to the high number involved in distinct behaviours between adult castes. Gender grouped more strongly than caste as expression was less variable between adult males than within each of the female castes. Similar findings have been presented for the social wasp *Vespula squamosa*, for which workers, queens and males clustered clearly into developmental stages (Hoffman and Goodisman, 2007). A study on the two fire ant species *Solenopsis invicta* and *S. richteri* also found expression patterns between developmental stages to differ more than between gender followed by caste and species (Ometto et al., 2011).

Our data confirmed, to some extent, previous findings for *B. terrestris* (Pereboom et al., 2005; Colgan et al., 2011). Several associations of gene functions with specific castes or developmental stages detected by Colgan et al. (2011) were also found in the present study. Discrepancies can be explained by, in contrast to our study, a lack of replication in the 2011 study or a difference in analysis structure; Colgan et al. (2011) implemented R-STAT (Stekel et al., 2000) to calculate differential expression of a contig within all libraries, whereas we performed specific pairwise comparisons. Little overlap could be found with an older study on caste determination in *B. terrestris* (Pereboom et al., 2005). However, due to the method implemented in that study, suppression subtractive hybridisation, only a few differentially expressed genes could be isolated, and also, due to a different focus, fewer comparisons were performed than in our study (Pereboom et al., 2005).

### Reproductive workers closely resemble queens

Towards the end of a bumble bee colony cycle a queen-worker conflict develops, in which reproductive workers compete with the mother queen for male parentage (Alaux et al., 2004; Bloch, 1999). The expression patterns observed in this study support our hypothesis that when bumble bee workers become reproductive they would, in comparison to highly eusocial species, more strongly resemble queens in their behaviour and physiology due to the more plastic nature of worker castes in bumble bees. Of all adult expression patterns, those of reproductive workers and mother queens were most similar, in fact more similar than between reproductive and non-reproductive workers. Only 111 transcripts differed significantly between reproductive workers and mother queens compared to 1,654 between reproductive and non-reproductive workers. Non-reproductive workers differed from mother queens even more strongly (5,316 DE transcripts). These findings are in strong contrast to patterns found in two highly eusocial hymenopteran species. In *A. mellifera* over 2,000 genes differed significantly in both comparisons between queens and either reproductive or non-reproductive workers; the expression of only 221 genes differed significantly between the two worker castes (Grozinger et al., 2007). Similarly, 2,785 genes were significantly up- or down-regulated between queens and reproductive workers in the myrmicine ant *Temnothorax longispinosus* compared to only 571 between reproductive and non-reproductive workers (Feldmeyer et al., 2014). Feldmeyer *et al*. (2014) suggested the high similarity between reproductive and non-reproductive workers in these two hymenopteran taxa indicates that a relatively low number of genes are required for ovary activation and egg laying compared to the high number involved in further physiological or behavioural differences which exist between queens and workers. Based on this assumption, our data indicate a greater similarity in behaviour and general physiology between bumble bee queens and reproductive workers than is the case for honey bees or myrmicine ants. The division of labour among bumble bees is not as clearly temporally or morphologically fixed as in the highly eusocial honey bees and most ants, indicating the capability of individual bumble bee workers to flexibly adapt their current role (e.g. from forager to nurse) to changing conditions within a colony at any given time (Cameron, 1989). In honey bees a shift towards a more ‘queen-like’ expression pattern was recorded in reproductive workers (Grozinger et al., 2007); but it is possible that the more flexible nature of the bumble bee worker roles in our study allowed a much stronger shift in behaviour and physiology, allowing the reproductive workers to more strongly resemble a queen.

### Male expression patterns are most distinct among adults

Males, in contrast to both queens and all workers, do not possess a sting and their antennae contain an additional segment. Their sexual organs naturally also differ. It was therefore surprising that the expression of comparatively few transcripts significantly differed during development. Within the larval stage no clear clusters could be formed based on expression patterns, and only 71 transcripts differed significantly in their expression levels between males and workers. During the pupal stage, when morphological features are being generated, expression patterns became more distinct with 162 transcripts differentially expressed. However, it was only in adulthood that the expression pattern of males became truly distinct from all other castes. In male adults between 2,411 and 8,706 transcripts were either up- or down-regulated compared to the three adult female castes mother queen, reproductive worker and non-reproductive worker. This indicates that a much greater number of genes may be required to control behaviour and the physiology of reproduction than to develop morphologies.

A high number (69; 59.5%) of the pupal transcripts up-regulated in male pupae, and therefore likely to contain some genes linked to the development of the male morphology, were of unknown function. The six *α-* and *β-tubulin* transcripts, which were over-represented in male pupae, are possibly linked to spermatogenesis as both *α2-* and *β-tubulin* are known to be testis specific in *Drosophila* (Theurkauf et al., 1986; Kemphues et al., 1979). 190 transcripts were involved in mitochondrial processes and a further 37 were associated with genes linked to muscle development. These 37 transcripts related to the proteins myosin, troponin, twitchin and titin, which are all integral parts of insect muscles (Hooper and Thuma, 2005). In their mating flights males have been recorded as covering significantly larger distances than workers from the same colony (Kraus et al., 2009). The apparent greater need for muscle development and higher energy levels in males compared to workers are possibly linked to their greater flight distances.

### Vitellogenin

Vitellogenin was originally thought to be limited to reproductive egg laying females due to its function as a yolk precursor in all oviparous animals, though it is now known to fulfil various functions in hymenopterans (Amdam et al., 2003). The reproductive ground plan model proposed by Amdam et al. (2004) describes how pleiotropic associations of reproductive genes, above all vitellogenin, with genes that control sensory perception, longevity and foraging behaviour have been utilised to control behaviour patterns in honey bee worker sub-castes.

Previously only one vitellogenin gene had been described for honey bees, which is differentially expressed in female castes (Amdam et al., 2012). However, in a more recent study on *Formica* ants four vitellogenin homologs were found within the genome of all ant and bee species included in the study (Morandin et al., 2014). These vitellogenin homologs were classed as conventional vitellogenin (*Vg-1*), *Vg-like-A*, *Vg-like-B* and *Vg-like-C*, which were expressed at different levels and differently between queens and workers. Four copies of *Vg-1* have been found in *Solenopsis invicta* (Wurm et al., 2011) and *Temnothorax longispinosus* (Feldmeyer et al., 2014) and two in *Pogonomyrmex barbatus* (Corona et al., 2013). In each of these cases, the gene copies showed differential expression between adult female castes.

Here we have found only one copy of *Vg-1* and two further vitellogenin genes which are closely related to *Vg-like-A* and *Vg-like-C*. *Vg-1*, as in *Formica* adults (Morandin et al., 2014), was the highest expressed of the three vitellogenin genes discovered in this study. We found *Vg-1* to be highly up-regulated in mother queens and reproductive workers compared to all other castes and developmental stages, which suggests it has maintained its conventional function in reproductive egg-laying females for *B. terrestris*. This also appeared to be the case for 3 out of 7 *Formica* species, in which *Vg-1* was up-regulated in queens compared to workers (Morandin et al., 2014). Workers were not grouped according to reproductive status in the Morandin *et al*. study (2014), which could explain the lack of significant differences between castes in more than 3 species. The comparison of expression between queens and workers for *Vg-A-like* differed among *Formica* species (up-regulated in queens for 3 and in workers for 1 species), (Morandin et al., 2014). In *B. terrestris* the homolog of *Vg-A-like*, XP_003400264, appears to play a lesser role in adults, as it was up-regulated in larvae and pupae of both genders compared to adults. Expression of *Vg-C-like* was significantly higher in workers than queens in all 7 *Formica* species (Morandin et al., 2014). In the current study the homolog of *Vg-C-like*, XP_003393940, was also down-regulated in mother queens but also in reproductive workers compared to higher levels in non-reproductive workers and adult males.

Here we have shown that three copies of vitellogenin genes are not only differentially expressed between adult females castes as shown for other hymenopteran taxa (Amdam et al., 2004; Morandin et al., 2014; Wurm et al., 2011; Feldmeyer et al., 2014; Corona et al., 2013), but that they are differentially expressed across all adult castes and between developmental stages.

### Carbohydrate processing enzymes

We found the expression of two carbohydrate processing enzymes to be differentially expressed among adult castes. Expression of *α-glucosidase* was almost exclusively restricted to female adults but with levels eight times higher in mother queens and reproductive workers than in non-reproductive workers. This is in contrast to honey bees for which *α-glucosidase* is down-regulated in reproductive compared to non-reproductive honey bee workers (Cardoen et al., 2011). In honey bees *α*-glucosidase catalyses the splitting of the sucrose present in nectar in the production of honey (Kubota et al., 2004; Ohashi et al., 1999). The apparent restriction of this protein to reproductive workers and mother queens may indicate a different role for this protein in *B. terrestris* compared to honey bees. *Glucose dehydrogenase*, on the other hand, was present in all *B. terrestris* adults but was down-regulated in reproductive workers and mother queens. The similar protein glucose oxidase is specifically found in the hypopharyngeal gland of forager honey bees and converts the glucose of nectar to gluconic acid and hydrogen peroxide in honey production (Ohashi et al., 1999). Glucose dehydrogenase may perform a similar function in *B. terrestris* as it also catalyses the oxidation of glucose to gluconic acid but without the by-product hydrogen peroxide (Bak, 1967). Expression of *α-glucosidase* significantly correlated positively with *Vg-1* while expression patterns of *glucose dehydrogenase* significantly correlated positively with *Vg-C-like*. These correlations indicate interactions between *vitellogenin* and the two carbohydrate enzymes, which may be associated with distinct foraging preferences among adult castes.

### Further caste specific genes

One highly represented gene in the list of transcripts over-expressed in mother queens compared to reproductive workers was serine protease inhibitor. Serine proteases have been detected in the venom of a variety of Hymenoptera species (Hoffman and Jacobson, 1996; Winningham et al., 2004). One possibility is that serine protease inhibitor was produced to counteract the effect of stings, either as a reaction to sting attacks or as a preventative measure. This could be linked to the high aggression shown towards a bumble bee queen by workers late in a colony cycle often resulting in her death (Bourke and Ratnieks, 2001).

Workers can become reproductive in queenright conditions, but whether workers or queens control worker reproduction is unresolved (Alaux et al., 2007). Intriguingly we found eight transcripts up-regulated in non-reproductive individuals (BTT06229 1, BTT09963 1, BTT20486 1, BTT15870 1, BTT22989 1, BTT27276 1, BTT17949 1 and BTT09790 1) whose expression is believed to be regulated by queen mandibular pheromone in *Apis mellifera* and where expression shows similar patterns (Grozinger et al., 2003; Cardoen et al., 2011). It is clear that further research is needed to understand the relationship between pheromonal signalling and ovary development (Amsalem et al., 2009).

In each of the caste comparisons performed in this study large numbers of differentially expressed transcripts either could not be associated with any known gene or were related to genes with so far unknown function. These range from 1,636 to 2,609 (32.0% - 54.4%) up-regulated transcripts when comparing between developmental stages. The number of differentially expressed transcripts was much lower between male and worker larvae (34 & 39) and pupae (128 & 34), but still the majority of these transcripts (58.7%) were of unknown function. 267 of the 791 fertility transcripts, i.e. up-regulated in reproductive workers and mother queens compared to non-reproductive workers, belonged to uncharacterised genes, while 465 and 526 transcripts in the comparison between non-reproductive workers and adult males were of unknown function. Clearly, further research is required in these areas.

## Conclusions

We conducted the first large scale RNA-seq analysis into caste differentiation within the genus *Bombus*, for which eusociality can be considered intermediate between that found in primitively eusocial taxa such as the paper wasp and highly eusocial species like the honey bee or most ants. As in other similar studies on eusocial hymenopterans, a high number of genes were differentially expressed in all comparisons between castes, genders and developmental stages. Significant overlaps with analyses on higher eusocial taxa exist in terms of overall expression patterns as well as specific genes. One striking difference between *B. terrestris* and higher eusocial hymenopterans is how much more closely a bumble bee reproductive worker resembles the queen regarding its gene expression. Further research may be able to determine whether this finding is restricted to *B. terrestris* or if it is linked to the more plastic nature of worker sub-castes in bumble bee taxa in general. The annotation of many unknown genes, which were differentially expressed in our analysis, and further research on *B. terrestris* following the imminent release of the genome will help us to better understand how distinct castes are created, maintained or altered within this important species.

## Acknowledgements

This study was funded by a NERC Biomolecular Analysis Facility research grant (NBAF 829). Illumina library preparation, sequencing and bioinformatics were carried out by Edinburgh Genomics, The University of Edinburgh. Edinburgh Genomics is partly supported through core grants from NERC (R8/H10/56), MRC (MR/K001744/1) and BBSRC (BB/J004243/1). M.C.H. is funded by a NERC PhD research grant. We would like to thank 3 reviewers and the editor for their useful comments on an earlier version of the paper.

## Data accessibility

All sequence data for this study are archived at European Genome-phenome Archive (EGA); accession number EGAS00001001169. Alignment files, raw read count lists, neighbor-joining tree files and the DESeq script are archived on Dryad (doi:10.5061/dryad.sp048). ANCOVA results, GO-analysis results and lists of differentially expressed transcripts are available as Supporting Information.

## Author contributions

All three authors developed the project idea, designed the experiment and were involved in the interpretation of data and finalisation of the manuscript. M.C.H. performed the experiment, analysed the data and drafted the manuscript.

